# Laboratory evolution in *Novosphingobium aromaticivorans* enables rapid catabolism of a model lignin-derived aromatic dimer

**DOI:** 10.1101/2024.03.22.586352

**Authors:** Marco N. Allemann, Ryo Kato, William G. Alexander, Richard J. Giannone, Naofumi Kamimura, Eiji Masai, Joshua K. Michener

## Abstract

Lignin contains a variety of interunit linkages, which leads to a range of potential decomposition products that can be used as carbon sources by microbes. β-O-4 linkages are the most common in native lignin and associated catabolic pathways have been well characterized. However, the fate of the mono-aromatic intermediates that result from β-O-4 dimer cleavage has not been fully elucidated. Here, we used experimental evolution to identify mutant strains of *Novosphingobium aromaticivorans* with improved catabolism of a model aromatic dimer containing a β-O-4 linkage, guaiacylglycerol-β-guaiacyl ether (GGE). We identified several parallel causal mutations, including a single nucleotide mutation in the promoter of an uncharacterized gene that roughly doubled the growth yield with GGE. We characterized the associated enzyme and demonstrated that it oxidizes an intermediate in GGE catabolism, β-hydroxypropiovanillone, to vanilloyl acetaldehyde. Identification of this enzyme and its key role in GGE catabolism furthers our understanding of catabolic pathways for lignin-derived aromatic compounds.

**Importance:** Lignin degradation is a key step for both carbon cycling in nature and biomass conversion to fuels and chemicals. Bacteria can catabolize lignin-derived aromatic compounds, but the complexity of lignin means that full mineralization requires numerous catabolic pathways and often results in slow growth. Using experimental evolution, we identified a new enzyme for catabolism of a lignin-derived aromatic monomer, β-hydroxypropiovanillone. A single mutation in the promoter of the associated gene significantly increased bacterial growth with either β-hydroxypropiovanillone or a related lignin-derived aromatic dimer. This work expands the repertoire of known aromatic catabolic genes and demonstrates that slow catabolism of lignin-derived aromatic compounds may be due to misregulation under laboratory conditions rather than inherent catabolic challenges.

## Introduction

Lignin is the second most abundant natural polymer and one of the three major components of plant cell walls (1). The lignin polymer consists of three main monomer units that differ based on the presence and number of methoxy units on the aromatic ring. In plants, these monomer building blocks are coupled via radical mechanisms to generate a wide variety of interunit linkages (2). Breakdown of lignin in the environment is thought to be performed mainly by fungi, which excrete extracellular peroxidases and laccases that cleave high molecular weight polymeric lignin into lower molecular weight aromatic compounds (3). These small soluble products can be further mineralized by other microbes. In bacteria, pathways have been described for conversion of diverse aromatic dimers into their constituent monomers, which can then be funneled into core catabolic pathways such as the various protocatechuate ring cleavage pathways (4, 5). These degradation pathways can also be used to valorize depolymerized lignin into value-added bioproducts (6, 7).

To date, pathways for catabolism of lignin-derived aromatic dimers connected by β-O-4 linkages have been the best characterized (Figure 1). This linkage typically comprises up to 40-50% of the inter-unit linkages found in lignin (1). A pathway for catabolism of dimers with β-O-4 linkages was first discovered over three decades ago in the alphaproteobacterium *Sphingobium lignivorans* SYK-6 (hereafter ‘SYK-6’) using the model compound guaiacylglycerol-β-guaiacyl ether (GGE) and has been extensively characterized at the genetic and biochemical level (8–14). More recently, additional enzymes for degradation of GGE and related intermediates have also been characterized in *Novosphingobium aromaticivorans* F199 (hereafter ‘F199’) (15–17) as well as other bacteria (18, 19). As shown in Figure 1, GGE catabolism in both F199 and SYK-6 proceeds via a set of conserved biochemical steps to produce the intermediates β-hydroxypropiovanillone (β-HPV) and guaiacol. In SYK-6, the β-HPV intermediate is funneled into the protocatechuate 4,5-cleavage pathway, and several of the necessary enzymes have recently been described (14, 20), while guaiacol is not assimilated. In contrast, previous work using F199 growing with GGE demonstrated that guaiacol was rapidly consumed but β-HPV accumulated transiently (15).

**Figure 1.**
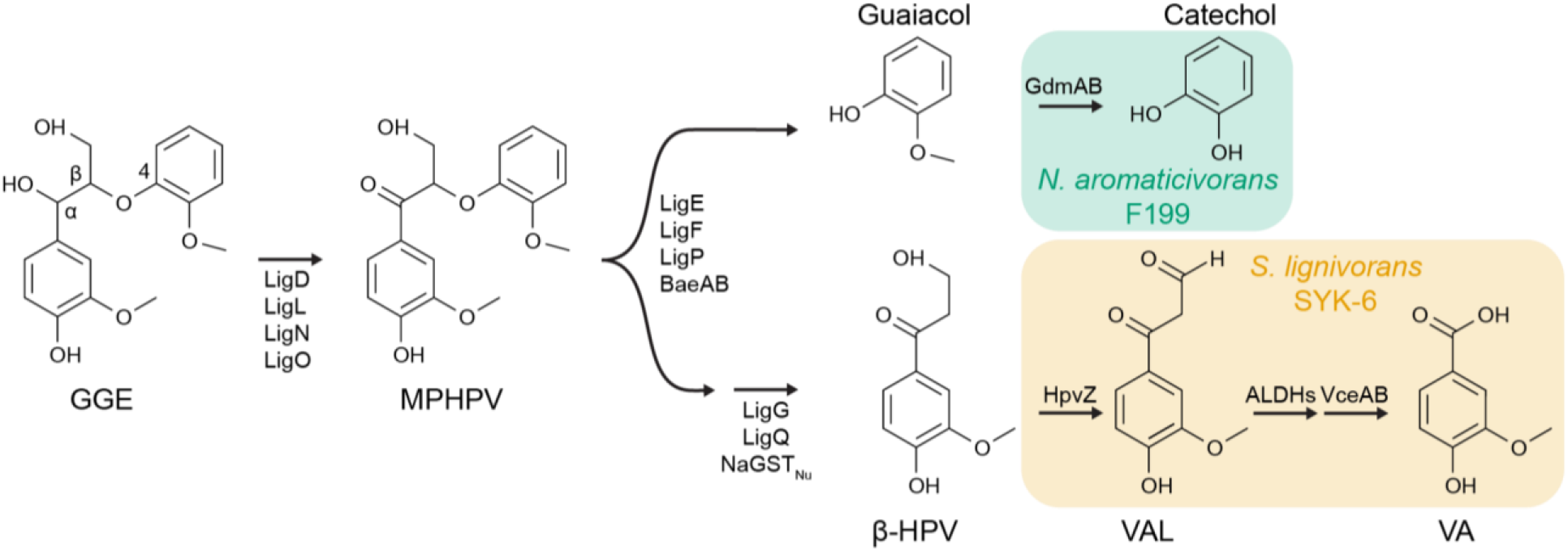
GGE catabolic pathways in two model lignin-degrading bacteria. The alcohol group at the α position of GGE is oxidized to form MPHPV. Various β-etherases cleave the β-ether group yielding β-HPV and, in the case of GGE, guaiacol. Previous works identified catabolic pathways for guaiacol in *N. aromaticivorans* F199 (17) and β-HPV in *S. lignivorans* SYK-6 (14, 20). For simplicity, stereochemistry of GGE and MPHPV is not shown. Abbreviations: GGE, guaiacylglycerol-β-guaiacyl ether; MPHPV, α-(2-methoxyphenoxy)-β-hydroxypropiovanillone; β-HPV, β-hydroxypropiovanillone; VAL: vanilloyl acetaldehyde; VA: vanillic acid. Enzymes: LigD, LigL, LigN, and LigO, short-chain dehydrogenase/reductases; LigE, LigF, LigP, BaeAB; β-etherases; LigG, LigQ, NaGST_Nu_ glutathione *S*-transferase; GdmAB, guaiacol *O*-demethylase; HpvZ, β-HPV oxidase; ALDHs, aldehyde dehydrogenases; VceA, VAA-converting enzyme; VceB, vanilloyl-CoA/syringoyl-CoA thioesterase

While both SYK-6 and F199 can grow with GGE as the sole carbon and energy source, neither wild-type strain grows rapidly under these conditions. We previously used experimental evolution to isolate a mutant of F199, which we termed JMN2, that grows more rapidly with GGE (21). Using barcoded transposon insertion sequencing in JMN2, we discovered a Rieske monooxygenase, GdmA, that demethylates guaiacol to catechol (17). Catechol can then be assimilated using a pathway for oxidative 1,2-cleavage of catechol that is natively present in F199. This newly-described pathway explains how F199 catabolizes the guaiacol produced from GGE degradation.

Since F199 was shown to accumulate but then fully catabolize β-HPV (15), we hypothesized that an additional pathway was present in this strain but inefficiently regulated under laboratory conditions. However, F199 does not contain a close homolog of the *hpvZ* gene that encodes the first enzyme for β-HPV catabolism in SYK-6 (20), so the identity of this potential catabolic pathway was unknown.

In this work, we describe further laboratory evolution experiments of F199 grown with GGE as the sole carbon and energy source. These evolution experiments yielded strains with a range of growth phenotypes, including one strain with a substantial improvement in yield compared to JMN2 during growth with GGE. Resequencing, mapping, and comparison of mutations across these evolved F199 strains led to the discovery of a novel β-HPV processing enzyme that we designate as HpvY. Heterologous expression in *E. coli* demonstrated that HpvY converts β-HPV to vanilloyl acetaldehyde (VAL).

## Materials and Methods

### Bacterial strains and growth conditions

*Escherichia coli* strains were routinely grown at 37°C in Lysogeny Broth (LB) unless stated otherwise. *Novosphingobium aromaticivorans* F199 strains were grown at 30 °C in R2A media (Teknova) unless noted otherwise. For growth of *N. aromaticivorans* strains in minimal media with defined carbon sources, DSM 457 media was used as described previously (24). For solid media, agar was included at 15 g/liter. The antibiotics kanamycin (50 μg/ml) and streptomycin (100 μg/ml) were used as required. Diaminopimelic acid (DAP) was added to LB as needed (60 μg/ml). All aromatic carbon substrates were dissolved in dimethylsulfoxide (DMSO) and added to minimal media for 1 g/L (w/v) final concentration. Due to the toxicity of guaiacol it was added to 0.25 g/L (2.0 mM).

### Laboratory evolution experiment

Triplicate cultures of wild-type *N. aromaticivorans* DSM12444 were inoculated from a single colony into DSM 457 with 2 g/L glucose and grown overnight at 30 °C. Cells were harvested by centrifugation and washed multiple times with DSM 457 lacking a carbon source and were inoculated into DSM 457 media supplemented with GGE (1 g/L) as a sole carbon source and incubated in a shaking incubator at 30 °C. Cultures were diluted 100-fold when growth was visually apparent by eye and significantly above the non-inoculated control. After approximately 2 months of growth and a total of five transfers, aliquots of the GGE grown cultures were streaked onto R2A agar for isolation of single colonies. Several purified colonies from each of the three evolutionary lineages were further screened for their abilities to grow with GGE and the best growing isolate derived from each lineage was chosen for further analysis.

### Resequencing and identification of mutations

High-quality genomic DNA was isolated from the evolved mutants using a Promega Wizard genomic DNA kit following manufacturer guidelines for Gram-negative bacteria with minor modifications. Genomic DNA was prepared for Illumina library construction and libraries were sequenced on the MiSeq Illumina platform to generate paired-end 300-bp reads using V3 chemistry. Sequence analysis and variant calling were performed in Geneious Prime (version 2021.0.3). Raw reads were trimmed with BBDuk plugin (version 1.0) with default settings. Trimmed reads were mapped to the *N. aromaticivorans* DSM1244 reference genome using the bowtie plugin (version 7.2.1). To identify single-nucleotide polymorphisms, the minimum variant frequency was set to 0.8. Mutations of interest were further verified by PCR and Sanger sequencing of purified products.

### Nanopore sequencing and genome assembly

10 ng of high-quality genomic DNA were amplified using the GenomiPhi v2 whole-genome amplification kit (Cytiva, Marlborough, MD), then the resulting DNA was simultaneously cleaned, concentrated, and size selected to remove fragments under 1 kbp by performing a bead cleanup with 0.4x volumes of HighPrep PCR cleanup beads (MagBio, Caithersburg, MD). 1.5 μg of this amplified DNA (hereto referred as wga DNA) was digested by 1.5 μL of T7 endonuclease (New England Biolabs, Ipswich, MA) in a 50 μL reaction for 1 h, then the digested wga DNA along with the native, unamplified DNA (hereto referred as nat DNA) was cleaned, concentrated, and size selected against fragments under 5 kbp using HighPrep PCR beads in a custom buffer containing 1.25 M NaCl and 20% PEG-8000 (25). The resulting DNA was used in conjunction with the Oxford Nanopore Technology (ONT, Oxford, UK) Rapid Barcoding Kit SQK-RBK004 to produce a sequencing library with nat and wga DNA barcoded separately, which was then loaded onto a MinION R9.4.1 flow cell driven by a Mk. IIB MinION device. The raw data were basecalled using the ONT Guppy basecaller on an HP Z8 workstation equipped with an NVIDIA Quadro RTX 4000 GPU with 8 GB of memory. The resulting reads for each sample underwent quality control using Filtlong v0.2.1 (https://github.com/rrwick/Filtlong) to remove fragments under 1 kbp, then to remove the bottom 5% of reads by quality. The resulting nat reads were used with Trycycler v0.5.1 (26) along with the assemblers Raven v1.5.3 (27), Miniasm + Minimap v0.3-r179 (28), and Flye v2.9 (29). This draft assembly was first polished using Medaka v1.5 (https://github.com/nanoporetech/medaka/) together with the nat reads, then Medaka polishing was repeated using the wga reads. Short-read polishing was then performed first with Polypolish v0.4.3 (30) then with Pilon v1.24 (31) to remove common errors associated with long-read assemblies (*e*.*g*. disagreement in homopolymer length).

### Targeted gene deletion mutagenesis

Chromosomal deletions in *N. aromaticivorans* were constructed using previously described methods with minor alterations (24). In-frame deletions were generated by allelic exchange using vector pAK405 (23). Briefly, homology arms containing 500-700 bp regions upstream and downstream of genes to be deleted were synthesized and cloned into pAK405 (Genscript, Piscataway, NJ). Constructs were mobilized into *N. aromaticivorans* via conjugation using the *E. coli* DAP auxotroph strain WM6026 with selection on R2A media containing kanamycin. Exconjugants were streaked to single colonies once from selection plates and grown overnight in R2A in the absence of kanamycin selection. Aliquots of the overnight culture were plated on R2A + streptomycin agar for counterselection against the integrated plasmid. Streptomycin-resistant colonies were patched to R2A + kanamycin + streptomycin and R2A streptomycin to screen for kanamycin sensitivity. Kanamycin-sensitive colonies were screened for gene deletions by colony PCR.

### Growth rate measurements

For the data shown in Figures 2, S1, S3, S5, and S6, strains were grown to saturation overnight in DSM 457 minimal media with 2 g/L glucose. Cells were pelleted and washed with DSM 457 lacking a carbon source before being diluted 100-fold into fresh medium containing the appropriate carbon source and grown as triplicate 1 mL cultures in 48-well plates (Greiner Bio-One, Kremsmünster, Austria). The growth was monitored by measuring the optical density at 600 nm (OD600) in an Epoch 2 plate reader (Agilent, Santa Clara, CA). The growth rates were calculated using the R package growthcurver (32).

**Figure 2.**
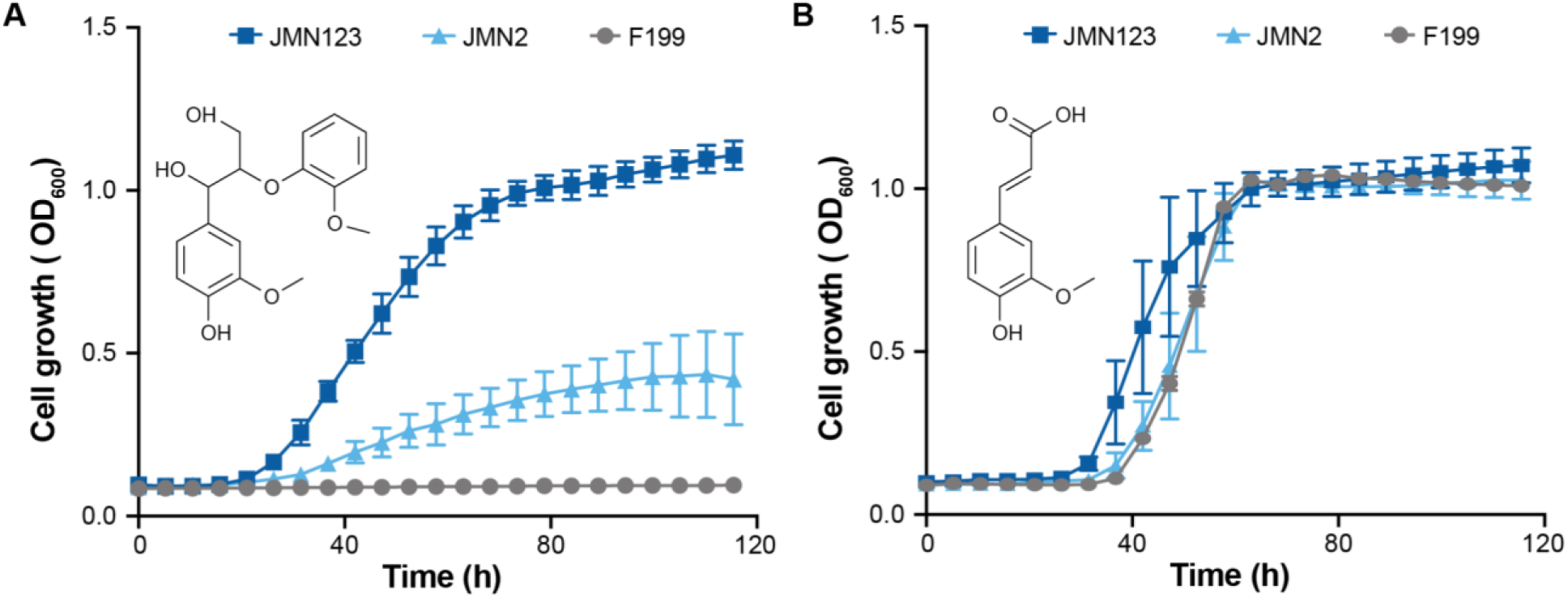
Adaptive laboratory evolution of *N. aromaticivorans* F199 for improved growth on GGE as a sole carbon source. (A) Growth of wild-type (F199) and evolved (JMN2, JMN123) strains with 1 g/L GGE (3.1 mM) as the sole carbon and energy source. (B) Growth of wild-type and evolved strains with 1 g/L ferulate (5.1 mM) as the sole carbon and energy source. Growth curves shown are the means of at least three independent experiments. Error bars show one standard deviation, calculated from the independent experiments.

For the data shown in Figure 3, strains F199, JMN2, JMN123, JMN159, and SYK-6 were cultured in LB for 24 h at 30°C. The cells were harvested by centrifugation at 14,000 × *g* for 1 min at 4 °C, washed twice with 3 mL of Wx medium, and then suspended in the same buffer. The cell suspensions were inoculated into Wx medium containing 2 mM GGE or HPV to an optical density at 600 nm (OD_600_) of 0.1 and incubated for 100 h at 30 °C with linear shaking at 567 cycles per minute using an EPOCH microplate reader (Agilent, Santa Clara, CA). The OD_600_ was measured continuously.

**Figure 3:**
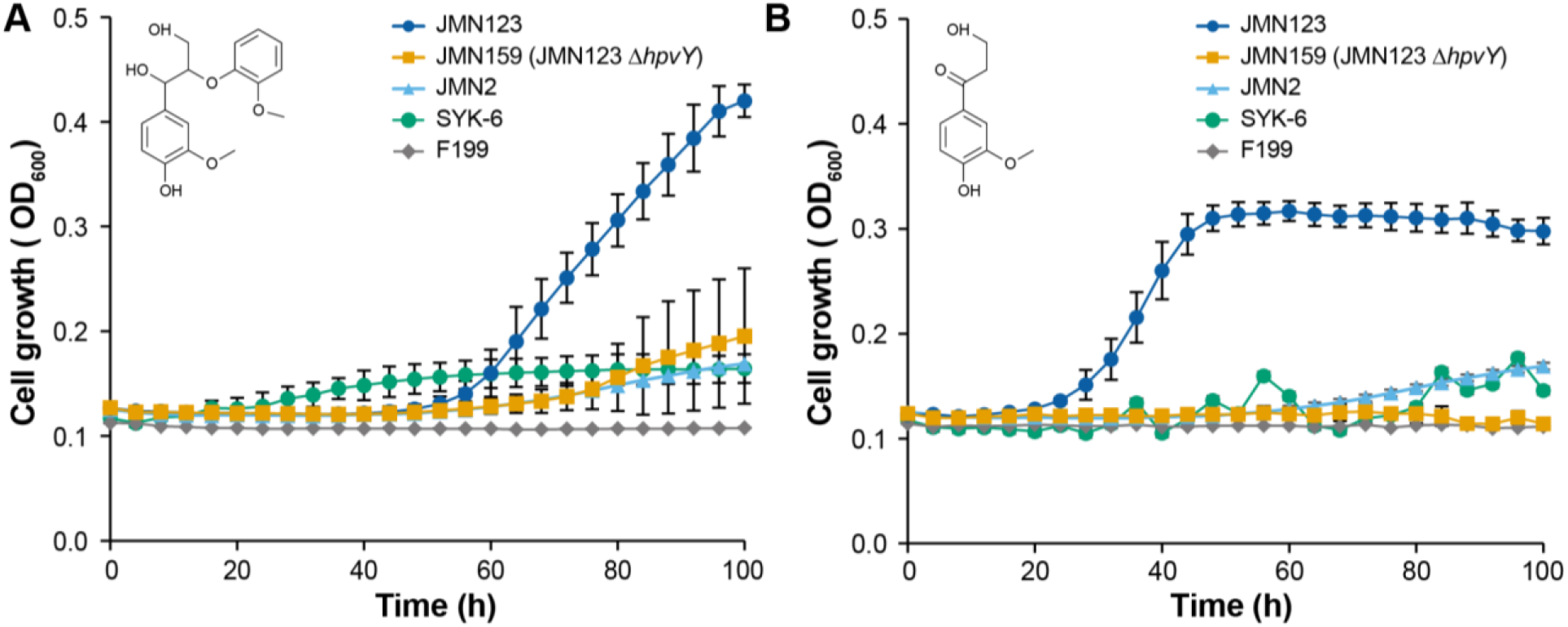
Impact of *hpvY* (*RS19130*) deletion during growth with GGE and β-HPV. (A) Deletion of *hpvY* from JMN123, yielding JMN159, significantly decreases growth yield with GGE. Strains were grown in minimal medium with 2 mM GGE as the sole carbon source. (B) Deletion of *hpvY* completely abolishes growth with β-HPV in JMN159. Strains were grown in minimal medium with 2 mM β-HPV as the sole carbon source. In both panels, error bars show the standard deviation calculated from three biological replicates.

### Metabolite accumulation assays

Strains were grown to saturation overnight in DSM 457 medium with 2 g/L glucose. Cells were harvested by centrifugation and washed several times with DSM 457 lacking a carbon source. Cells were then diluted into fresh DSM 457 medium containing 1 g/L GGE and incubated at 30 °C. At the indicated times ∼1 mL aliquots of the culture were removed and cells pelleted and supernatant was collected and filtered to 0.22 µm. Filtered supernatant samples were then analyzed by LC-MS to monitor intermediate product accumulation across the strains over time. For each sample, 5 µL of supernatant was split-loaded onto an in-house pulled nanospray emitter (75 µm inner diameter) packed with 15 cm of C18 resin (1.7 µm Kinetex; Phenomenex) and separated over a 15 min reversed-phase gradient using a Vanquish HPLC interfaced directly to a Q Exactive Plus mass spectrometer (Thermo Scientific) (21). Eluting analytes were measured by the Q Exactive operating in negative ion mode monitoring a mass-to-charge range of 100-500 *m*/*z*; resolution 35,000; 3 microscan spectrum averaging). Peak areas were extracted for MPHPV (317.1031 [M-H]) and β-HPV (195.0663 [M-H]) pathway intermediates via Skyline software (33) and areas compared across samples to assess strain bottlenecks.

### Preparation of resting cells

Cells of *E. coli* transformants, F199, JMN2, JMN123, and JMN159 were grown in LB for 12 h (*E. coli* strains) and 24 h (F199 and F199-derivative strains) at 30°C with linear shaking at 160 rpm. F199 derivatives were cultured in Wx medium containing 10 mM sucrose, 10 mM glutamate, 0.13 mM methionine, and 10 mM proline (Wx-SEMP) and Wx-SEMP containing 2 mM GGE, 2 mM HPV, or 2 mM Guaiacol for 12 h at 30°C. The cells were collected by centrifugation (14,000 ’ *g* for 1 min), washed twice with 50 mM Tris-HCl buffer (pH 7.5), resuspended in the same buffer, and used as resting cells.

### Conversion of b-HPV by F199, F199-derivative strains, and *E. coli* expressing *hpvY*

Resting cells of F199, JMN2, JMN123, JMN159, and *E. coli* BL21(DE3) harboring p16hpvY (F199 and F199-derivative strains, OD_600_ of 5.0; *E. coli* transformant, OD_600_ of 10) were incubated with 100 µM HPV at 30°C with shaking for 8–24 h. The supernatant obtained by centrifugation at 19,000 × *g* for 1 min at 4°C was analyzed by HPLC.

### Expression of *hpvY* in *E. coli*

A DNA fragment carrying *hpvY* (RS19310) was amplified through PCR using the F199 total DNA and the primer pairs listed in Table 2. The amplified fragment of *hpvY* was cloned into pET-16b using an NEBuilder HiFi DNA assembly cloning kit to form p16hpvY. *E. coli* BL21(DE3) cells harboring p16hpvY were grown in LB, and gene expression was induced for 4 h at 30°C by adding 1 mM isopropyl-b-D-thiogalactopyranoside when the OD_600_ of the culture reached 0.5. Gene expression was examined using sodium dodecyl sulfate–12% polyacrylamide gel electrophoresis. The protein bands in gels were stained with Coomassie brilliant blue.

**Table 1.**
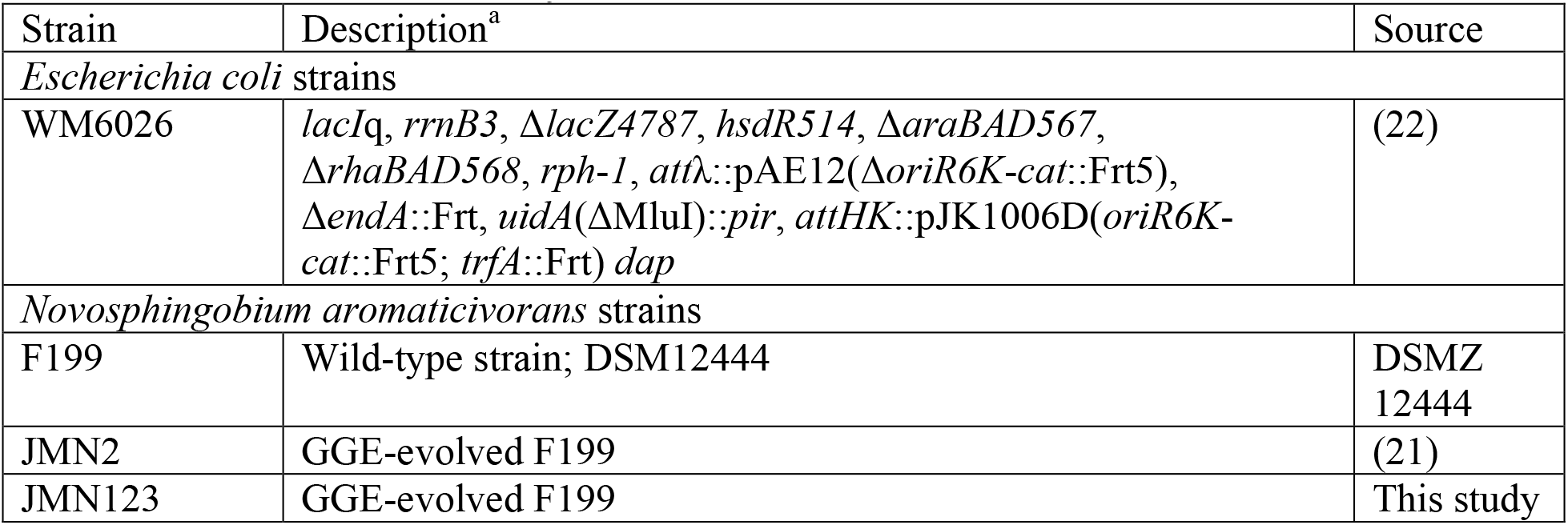
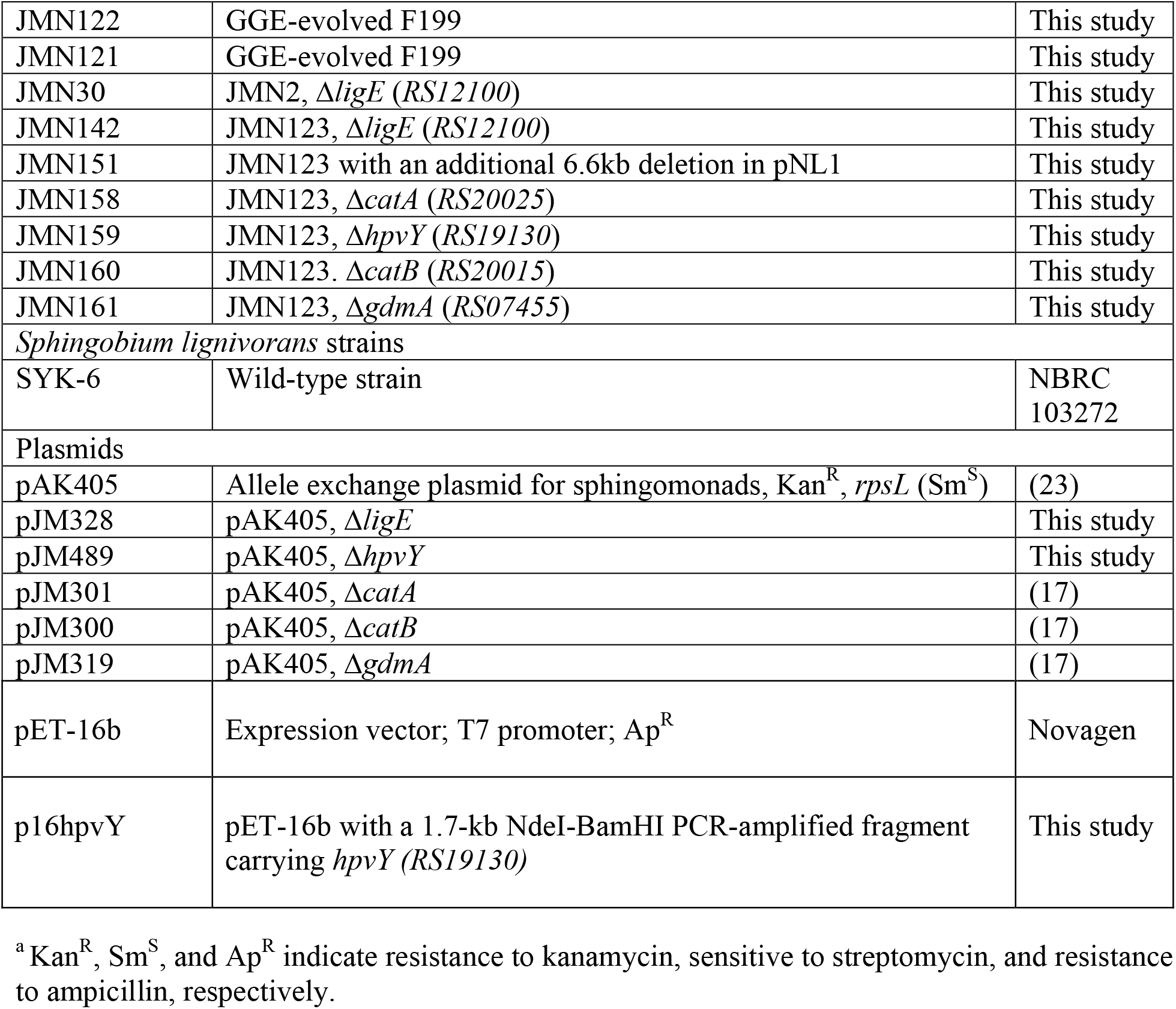
Strains used in this study.

| Strain | Description<sup>a</sup> | Source |
| --- | --- | --- |
| *Escherichia coli* strains |  |  |
| WM6026 | *lacIq*, *rrnB3*, Δ*lacZ4787*, *hsdR514*, Δ*araBAD567*, Δ*rhaBAD568*, *rph-1*, *attλ::pAE12*(Δ*oriR6K-cat::Frt5*), Δ*endA::Frt*, *uidA*(Δ*MluI*)::*pir*, *attHK::pJK1006D*(*oriR6K-cat::Frt5*; *trfA::Frt*) *dap* | (22) |
| *Novosphingobium aromaticivorans* strains |  |  |
| F199 | Wild-type strain; DSM12444 | DSMZ 12444 |
| JMN2 | GGE-evolved F199 | (21) |
| JMN123 | GGE-evolved F199 | This study |
| JMN122 | GGE-evolved F199 | This study |
| JMN121 | GGE-evolved F199 | This study |
| JMN30 | JMN2, $\Delta ligE$ (RS12100) | This study |
| JMN142 | JMN123, $\Delta ligE$ (RS12100) | This study |
| JMN151 | JMN123 with an additional 6.6kb deletion in pNL1 | This study |
| JMN158 | JMN123, $\Delta catA$ (RS20025) | This study |
| JMN159 | JMN123, $\Delta hpvY$ (RS19130) | This study |
| JMN160 | JMN123, $\Delta catB$ (RS20015) | This study |
| JMN161 | JMN123, $\Delta gdmA$ (RS07455) | This study |
| *Sphingobium lignivorans* strains |  |  |
| SYK-6 | Wild-type strain | NBRC 103272 |
| Plasmids |  |  |
| pAK405 | Allele exchange plasmid for sphingomonads, Kan<sup>R</sup>, *rpsL* (Sm<sup>S</sup>) | (23) |
| pJM328 | pAK405, $\Delta ligE$ | This study |
| pJM489 | pAK405, $\Delta hpvY$ | This study |
| pJM301 | pAK405, $\Delta catA$ | (17) |
| pJM300 | pAK405, $\Delta catB$ | (17) |
| pJM319 | pAK405, $\Delta gdmA$ | (17) |
| pET-16b | Expression vector; T7 promoter; Ap<sup>R</sup> | Novagen |
| p16hpvY | pET-16b with a 1.7-kb NdeI-BamHI PCR-amplified fragment carrying *hpvY* (RS19130) | This study |

<sup>a</sup> Kan<sup>R</sup>, Sm<sup>S</sup>, and Ap<sup>R</sup> indicate resistance to kanamycin, sensitive to streptomycin, and resistance to ampicillin, respectively.

**Table 2.**
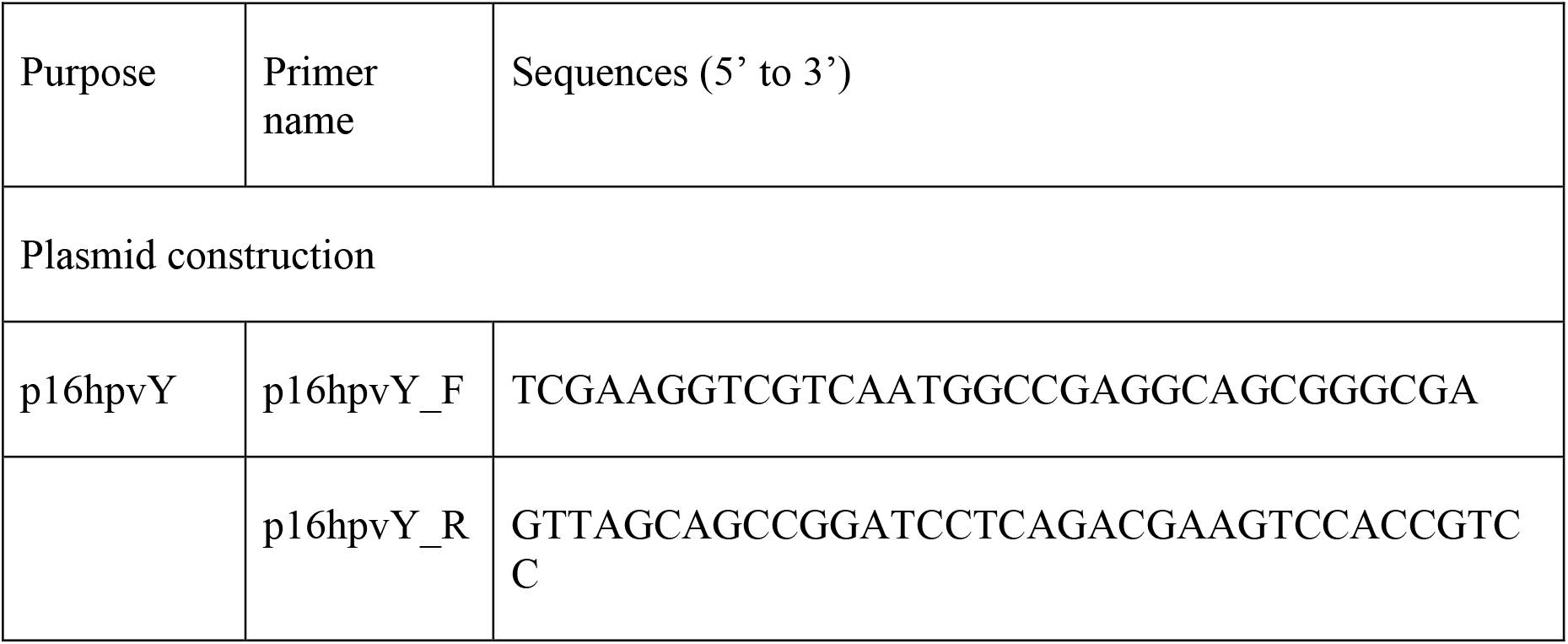
primers used in this study.

### HPLC analysis

HPLC analysis was conducted using the ACQUITY UPLC system (Waters). The sample was filtered through a PTFE filter (Captiva Econofilter, Agilent) with a pore size of 0.20 µm and then analyzed using a TSKgel ODS-140HPT column (particle size, 2.3 µm; 2.1 x 100 mm, Tosoh). Analysis of HPV, VAL-Tris, and VAA was performed in gradient mode. The mobile phase was a mixture of solution A (acetonitrile containing 0.1% formic acid) and B (water containing 0.1% formic acid) under the following conditions: 0-3.2 min, 5% A; 3.2-6.0 min, linear gradient from 5.0 to 40% A; 6.0-6.5 min, decreasing gradient from 40 to 5.0% A; 6.5-7.0 min, 5% A. The flow rate was 0.5 mL/min, and the column temperature was 30 °C. HPV, VAL-Tris, and VAA were detected at 310 nm.

## Results and Discussion

### Laboratory evolution of F199 for growth with lignin-derived aromatics

We previously described the evolution and resequencing of strain JMN2, a mutant of F199 that was selected for growth with GGE as the sole carbon source (17, 21). To identify other potential mutations that improve growth with GGE, we initiated additional evolution experiments. Three independent cultures of F199 were grown and serially passaged in minimal media containing GGE as the sole carbon source. After approximately 33 generations, each mixed evolution culture showed improved growth rate and yield compared to the F199 parent. We then isolated approximately eight single colonies from each replicate culture and assayed growth with GGE as the sole carbon and energy source. Each culture yielded at least one isolate that showed significantly improved growth with GGE, and the top-performing strain from each replicate culture was selected and preserved for further analysis (Figure 2A and Figure S1). The growth improvements in evolved strains were specific to GGE and not observed with other lignin-derived aromatic compounds such as ferulate (Figure 2B).

### Identification of mutations and deletions in evolved lineages

To gain insight into the genetic basis for the evolved phenotypes, we resequenced the genomes of all three isolates using short reads and, to detect any large scale rearrangements, generated a *de novo* genome sequence of JMN123 using Nanopore sequencing. No large rearrangements were identified, and a full comparison of the various point mutations found in these strains is shown in Supplementary Table 1. All four strains contain parallel mutations, including large deletions in plasmid pNL1 and mutations upstream of *Saro_RS12100*.

### Role of *ligE* in GGE metabolism

Upon examination and comparison of the mutations found within the four evolved strains, we noted that all four contain an identical point mutation 55 bp upstream of the start codon of *Saro_RS12100*. This gene encodes for LigE, one of the β-etherases known to be involved in GGE catabolism (10, 13). Based on sequence analysis, this upstream region contains plausible - 35 and -10 promoter sites as well as an inverted repeat motif (Figure S2). The observed point mutation occurs in the predicted inverted repeat motif and could potentially interfere with binding of a transcriptional regulator (34). Repeated attempts to reconstruct this single nucleotide mutation in a wild-type background were unsuccessful. Instead, an in-frame unmarked deletion of *ligE* was engineered in both JMN2 and JMN123 and the resulting strains were assessed for their growth in minimal medium with GGE. In both genetic backgrounds, the *ΔligE* mutation severely hindered growth with GGE (Figure S3). Given the phenotypes observed, we propose that the *ligE* promoter mutation was most likely a gain of function mutation that increases *ligE* expression and therefore improves GGE conversion to monomers.

### Effects of deletions in pNL1

Wild-type F199 contains a large plasmid, pNL1, with many accessory catabolic genes (35). Based on previous sequencing and genetic characterization of pNL1, this plasmid can be divided into three distinct regions based on predicted gene functions within each region: replication, mobilization, and aromatic degradation. All four evolved strains contained significant deletions in the aromatic degradation region, which contains a catechol 2,3 cleavage pathway (*xylEGHIJKQ*) (Figure S4). This *xyl* pathway was previously shown to contribute to the degradation of protocatechuate (PCA) in a previous study focused on aromatic pathway discovery in F199 (24).

An additional 6.6 kb of the aromatic degradation region was deleted in JMN2 than in JMN123. We hypothesized that this genetic difference could explain the improved yield of JMN123 during growth with GGE. To test this hypothesis, we made an additional targeted deletion in pNL1 of JMN123 to replicate the deletion seen in JMN2 (Figure S5A), yielding strain JMN151. JMN151 had a slightly lower yield than JMN123 during growth with GGE but was still substantially improved compared to JMN2 (Figure S5B). We conclude that differences in the extent of deletion of pNL1 may affect growth with GGE but are not the major contributor to the improved growth by strain JMN123.

### Identification of a new enzyme and characterization of its role in GGE catabolism

Further comparison of the mutations found in the four evolved isolates identified a single nucleotide mutation that was uniquely found in JMN123, 49 bp upstream of the start codon of *Saro_RS19130*. As with the *ligE* mutation, this mutation is found in a region upstream of the coding sequence of *RS19130* that contains a predicted promoter (Figure S6). This gene encodes a predicted choline dehydrogenase that has 39% amino acid sequence identity to HpvZ from SYK-6. Based on this homology, we refer to *RS19130* as *hpvY*. An in-frame deletion of *hpvY* was constructed in JMN123 to yield strain JMN159. When strains JMN2, JMN123, and JMN159 were grown with GGE as a sole carbon source, JMN159 showed a significant decrease in yield, comparable to JMN2 (Figure 3A).

To further identify the role of *hpvY* in GGE catabolism, we measured growth of the evolved and engineered strains with β-HPV as the sole carbon and energy source (Figure 3B). JMN123 grew rapidly with β-HPV, while deletion of *hpvY* in JMN159 completely abolished growth with β-HPV. The growth defect in JMN159 was specific to GGE and β-HPV, and was not observed with a related lignin-derived aromatic compound, ferulate (Figure S7). We conclude that the mutation in JMN123 upstream of *hpvY* activated a previously-unidentified pathway for β-HPV catabolism.

We also monitored metabolite accumulation during growth with GGE to identify pathway bottlenecks that were introduced or removed by alterations to *hpvY*. Similar to the previous experiments, we grew JMN2, JMN123, and JMN159 with GGE as the sole source of carbon and energy and monitored the concentration of intermediate metabolites MPHPV and β-HPV (Figure 4). JMN2 accumulated significant concentrations of both intermediates, while JMN123 accumulated these intermediates only transiently and then fully consumed them. Deletion of *hpvY* in JMN159 again led to accumulation of these intermediates, consistent with a role for HpvY in β-HPV catabolism. We hypothesize that accumulated β-HPV inhibited the glutathione transferases responsible for conversion of MPHPV to β-HPV and resulted in MPHPV accumulation.

**Figure 4:**
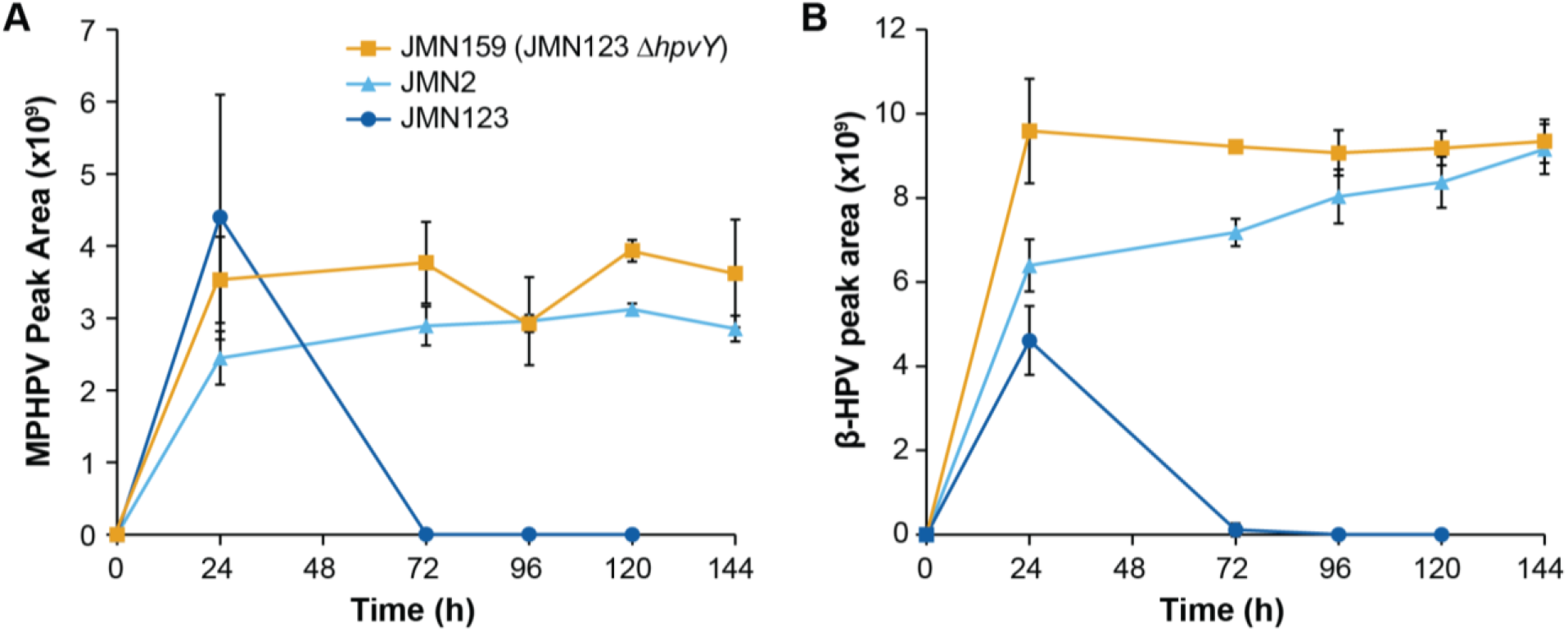
Impact of *hpvY* (*RS19130*) deletion on GGE conversion. Cells were grown in minimal medium containing 1 g/L GGE (3.1 mM) as the sole carbon source. Accumulation of (A) MPHPV and (B) β-HPV were monitored by LC-MS. Error bars show one standard deviation, calculated from three biological replicates.

### Role of HpvY in β-HPV catabolism

Low accumulation of β-HPV during growth with GGE suggested that JMN123 has higher β-HPV conversion activity than the other *N. aromaticivorans* strains tested. To more directly measure β-HPV conversion and to test the regulation of this activity, we grew strains of *N. aromaticivorans* with four different media, harvested the cells, added β-HPV, and measured the disappearance of this substrate by HPLC. When cells were grown in Wx-SEMP medium, minimal β-HPV conversion activity was detected (Figure 5A). Pre-culture with 2 mM GGE strongly induced β-HPV conversion by JMN123 but not by JMN159 (JMN123 Δ*hpvY*) (Figure 5B). Pre-culture with lysogeny broth (LB) or Wx-SEMP + 2 mM guaiacol also induced β-HPV conversion activity (Figure S8). We conclude that *hpvY* is required for β-HPV conversion and that this pathway is inducible in JMN123, either by metabolites in the GGE catabolic pathway or by a constituent of complex media. The specific inducer molecule and putative transcriptional regulator have not yet been described.

**Figure 5:**
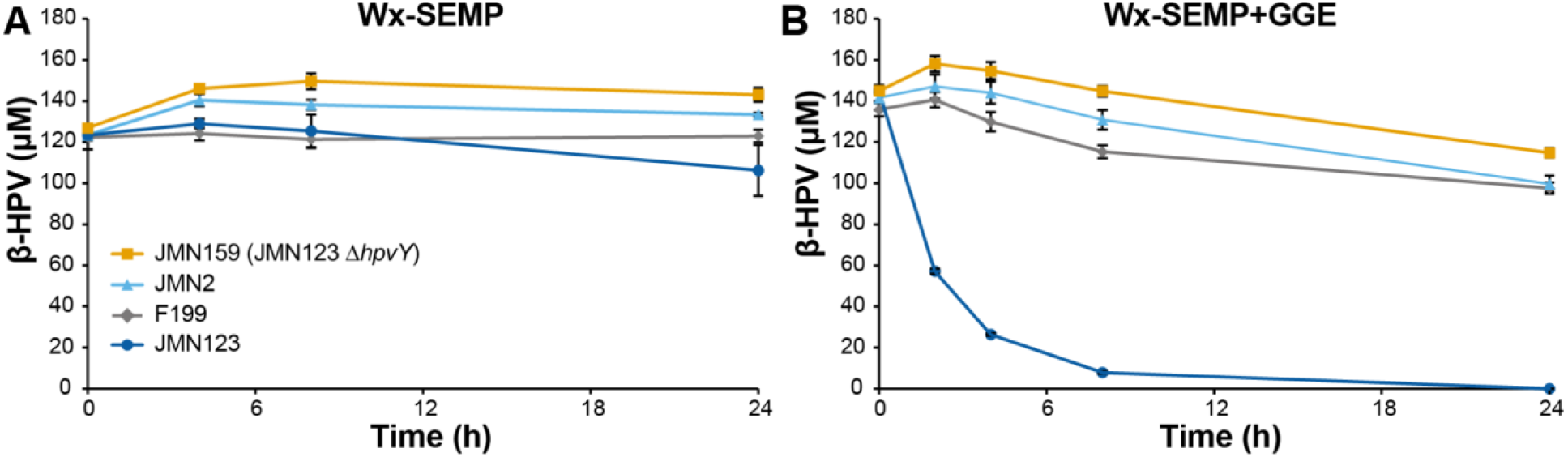
Conversion of β-HPV by resting cells of *N. aromaticivorans*. Strains were pre-cultured in (A) Wx-SEMP or (B) Wx-SEMP + 2 mM GGE, concentrated, and incubated with β-HPV. Residual β-HPV was detected by HPLC. Error bars show the standard deviation, calculated from three biological replicates.

### Heterologous expression of HpvY and demonstration of β-HPV conversion

While β-HPV conversion by JMN123 requires *hpvY*, the effect of HpvY on β-HPV conversion could be indirect. To identify the activity of HpvY, we heterologously expressed *hpvY* in *Escherichia coli* BL21(DE3). We observed expression of a protein with the expected mass of approximately 60 kDa in both the soluble and insoluble fractions (Figure S9). We therefore measured conversion of β-HPV using resting cells of control and *hpvY*-expressing *E. coli*. No conversion of β-HPV was observed by the control cells, while the cells expressing HpvY produced vanilloyl acetaldehyde (VAL) and vanilloyl acetic acid (VAA) (Figure 6). VAA is likely produced through promiscuous oxidation of VAL by endogenous aldehyde dehydrogenases (Figure 1). Based on these results, we conclude that HpvY directly oxidizes β-HPV to VAL, equivalent to the reaction catalyzed by HpvZ (20). A precise comparison of catalytic properties between HpvY and HpvZ would require additional enzyme purification.

**Figure 6:**
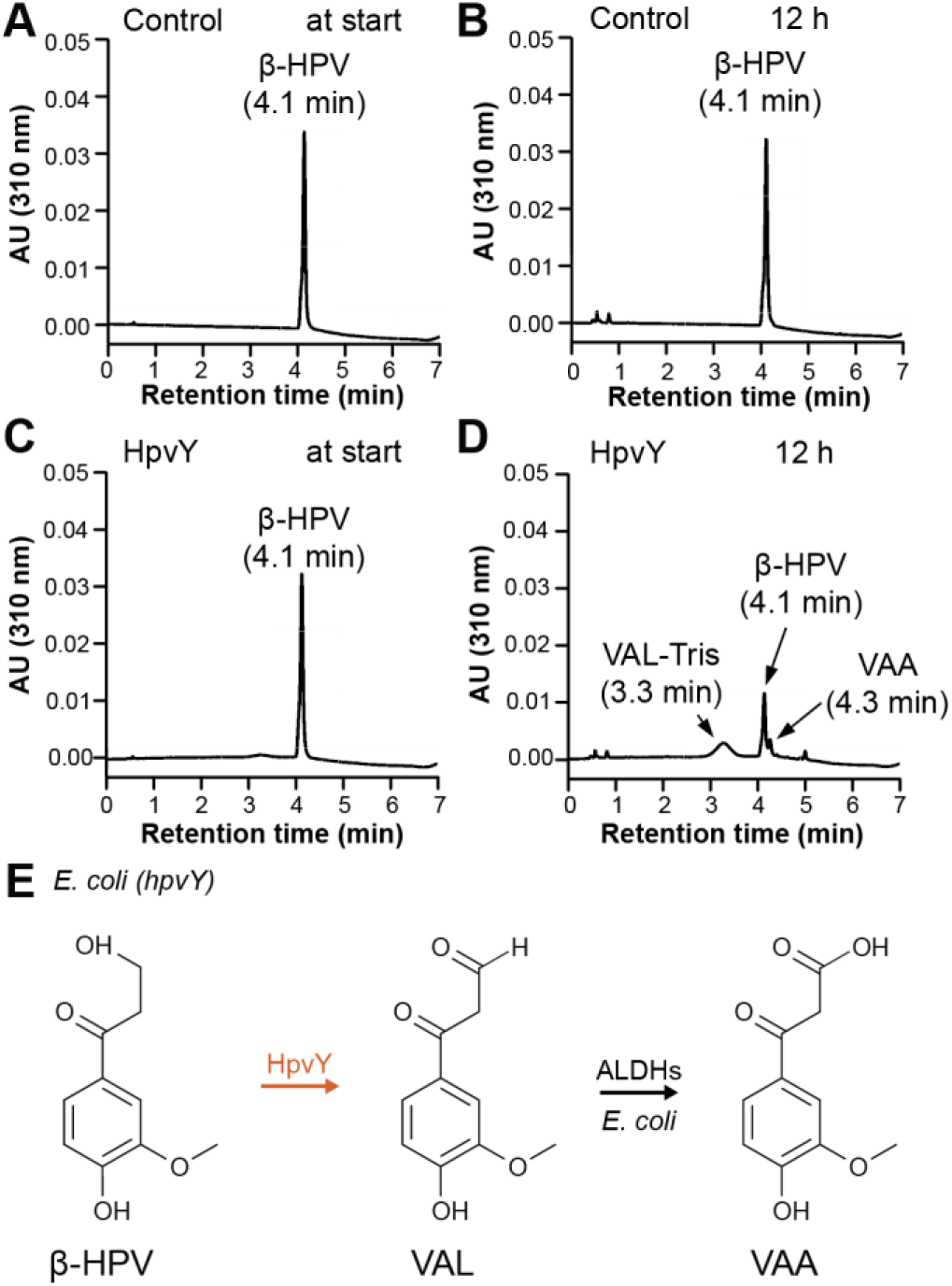
HPLC analysis of β-HPV conversion by resting *E. coli* cell suspensions containing either an empty plasmid control (A+B) or a plasmid heterologously expressing *hpvY* (C+D). 100 μM β-HPV was added at t_0_. β-HPV, VAA, and VAL-Tris are identified by comparison to authentic standards. (E) Proposed reaction scheme in *E. coli* heterologously expressing HpvY. Abbreviations: β-hydroxypropiovanillone, β-HPV; VAL, vanilloyl acetaldehyde; VAL-Tris, imine derivative of VAL produced non-enzymatically by reaction with Tris (20); VAA, vanilloyl acetic acid.

Similar to SYK-6 and resting *E. coli*, it is likely that VAL produced by HpvY in F199 is oxidized to VAA. In SYK-6, VAA is then converted to VA by the actions of VceA, VceB, and an unidentified enzyme. However, homologs of VceA and VceB are not found in F199, suggesting that additional pathway enzymes remain to be discovered.

## Conclusion

In this work, we used adaptive laboratory evolution to generate a mutant strain of *N. aromaticivorans* F199 that rapidly and completely catabolizes a model lignin dimer. Resequencing multiple evolved strains identified several key parallel mutations that contribute to the improved growth phenotype. Further analysis of these mutations led to the discovery of a novel β-HPV converting enzyme, designated HpvY, that converts β-HPV into VAL. This work highlights the utility of experimental evolution for both pathway optimization and pathway discovery.

## Acknowledgements

This manuscript has been authored by UT-Battelle, LLC under Contract No. DE-AC05-00OR22725 with the U.S. Department of Energy (DOE). This work was primarily supported by the U.S. DOE, Office of Science, Office of Biological and Environmental Research, though an Early Career Award to JKM. Metabolite analysis by RJG was supported by the Center for Bioenergy Innovation (CBI), U.S. Department of Energy, Office of Science, Biological and Environmental Research Program under Award Number ERKP886. Analysis of HpvY was supported by a JST grant JPMJPF2104. This work also used the resources of the Compute and Data Environment for Science (CADES) at Oak Ridge National Laboratory, which is supported by the U.S. Department of Energy’s Office of Science under Contract No. DE-AC05-00OR22725. The authors wish to thank Leah Burdick for assistance with genomic DNA sequencing.

